# Diversified Learning Environment Alleviates Learning Impairment Caused by Contextual Stress

**DOI:** 10.1101/2020.01.15.907840

**Authors:** Chen Gong, Meng Zhang, Yanjie Zhang, Xiang Liu, Pan Liu, Xiaoling Liao, Yi Zhou, Zhiyue Shi, Xue Liu

**Author notes:** Correspondence should be addressed to: Zhiyue Shi or Xue Liu.

## Abstract

Environment is capable of affecting our learning ability. Existing studies reported that learning ability can be enhanced by enriched environment and impaired by stressful context. However, it is still unclear whether diversified environment can reverse or ameliorate the learning difficulty caused by stressful context. In this study, several behavioral tasks were designed to test the role of diversified environment in active avoidance learning. In the present study, sound-cued active avoidance (two-way shuttle box) acted as learning paradigm. Multiple shuttle boxes with the identical size but different designs were employed to mimic diversified environment in learning tasks. Mild but inevitable foot shocking was adopted to increase animal’s stress to certain context. To quantify the depression/anxiety level of animals, open field test, forced swimming test, light-dark box test, and elevated plus maze were performed. The following findings were reported. First, diversified learning environment could improve learning ability in active avoidance, as manifested by higher successful rate and sharper learning curve. Second, elevating the stress level of animal to a certain context could noticeably reduce its performance in active avoidance learning. Third, the learning impairment attributed to stressful context can be improved by training in diversified environments. Thus, as revealed from the results, learning impairment caused by stressful context can be alleviated by diversified learning environment which may facilitate further medical and education applications.

## Introduction

Environmental stress significantly impacts learning ability of both human and mammalians[1]. However, stress will exert different or adverse effects on learning performance. As reported in previous research, appropriate stress stimuli during training can elevate learning scores[2]. In human studies, an individual was exposed to stress for a relatively long time before learning task could promote learning[3, 4]. In rodents, rats trained in cold water maze (19°C, i.e., under environmental stress) achieved better learning results than those trained in warm water maze (25°C)[5]. Besides, under stressful environment, the learning ability of both humans and animals would be impaired, which is a major cause of depression or post-traumatic stress disorder[6]. In humans, stress induced learning deficits are considered one major symptom attributed to negative emotions [7, 8]. Numerous existing studies also reported that acute social or physical stress could severely impair memory recovery [9-11].

Two different approaches have been developed to counter the adverse impacts of environmental stress. One approach is known as the enriched living environment that has been proved as an effective way to alleviate animal stress and improve learning ability by many references[12-14]. In rats, a commonly used method to enrich the environment is to keep the subject in a large space filled with daily renewed objects and socially contact with the identical group [15, 16]. Rats bred in enriched environments exhibit better learning and memorizing abilities than isolated littermates [17-19]. Several studies further reported that the simple behavioral therapy in enriched environment could effectively improve neural function and mitigate age-related memory dysfunction in rats and mice [13]. In addition, enriched environment could be critical to reduce negative emotions in animals [20]. It was reported that enriched environment could mitigate the enhancing effect of addictive drugs and exert similar effects to anti-depressants [21] [22]. Furthermore, environmental enrichment has been proposed as a “therapeutic” model in which rats are bred in an isolated or enriched environment after being exposed to drugs or stress [23, 24].

The other approach is to diversify the training environment. Exchanged students between schools, cross department training, or field training at different places for professional athletes have been widely used and proved effective to enhance performance in numerous aspects. However, the role of diversified environment in learning still remains to be elucidated. Two-way shuttle box acts as a feasible behavioral paradigm to study active avoidance, and it has been employed to investigate the relationships between stress in training environment and animal learning ability[25]. In active avoidance, animal’s behavior is related to the level of fear or stress to neutral conditional stimulus[26]. Meanwhile, excessive environmental stress can adversely affect behavior performance in active avoidance task. Since the training and testing of this paradigm have been commonly performed in the identical shuttle box, when the animal is progressively learning to respond to conditioned stimuli, it may develop more fear to the environment (shuttle box) [25, 27-29]. Such environmental stress is likely to interfere with and down-regulate the success rate of the animal’s active avoidance behavior. In other words, when an animal perceives a conditional stimulus, it is supposed to balance the conditional punishment with environmental punishment before making a decision[26, 29, 30].

The present study hypothesized that diversified training environments can improve learning ability by lowering the negative effects caused by environmental stress. Multiple two-way shuttle boxes were adopted to verify whether diversified training environments can improve the animal’s learning ability. Three shuttle boxes with same size but slightly different designs were adopted, and animal’s performance to conditional sound stimuli in individual or combined boxes was recorded. Moreover, based on multiple standards (e.g., open field test, forced swimming test, light-dark box test, as well as elevated plus maze), mental status of animals was also quantified. It is found that animals trained in diversified environments showed better learning abilities. Increased environmental stress by unconditioned foot shock evidently impair animal’s active avoidance learning. Note that impaired learning ability attributed to environmental stress can be relieved by diversified training environments. Our results suggested that stressful context can harm the learning in active avoidance tasks and this impairment can be alleviated by simple but diversified environment.

## Materials and Methods

### Animals

Adult Sprague–Dawley rats (female, 2 months, 180–210 g) were purchased from the Laboratory Animal Center at the Third Military Medical University. All experimental procedures were performed in accordance with institutional animal welfare guidelines and approved by the Third Military Medical University Animal Care and Use Committee. In this study, animals (adult rats) were maintained on a 12-h light/dark cycle with free access to food and water at a room temperature of 25–28°C. All efforts were made to minimize animal suffering. Minimal number of animals required for statistical reliability were recruited. Each experimental process approved by the Army Military Medical University Ethical Committee for Animal Research (SYXK-PLA-20120031) was performed following the National Institutes of Health Guide for the Care and Use of Laboratory Animals.

### Behavioral Procedures

#### Sound-cued active avoidance training

The experiment box was split into two chambers. The opening between two chambers was 10 cm. The bottoms of the two chambers were both covered with a stainless-steel grid that could be powered by direct current (0.5 mA). The animals were placed in a shuttle box and subsequently programmed to get acclimated for five minutes without being stimulated. After the pre-operation was completed, the sound was played as the stimulation for 6s. After the playback conditional stimulation was performed, the system recognized the position of the animal within 0.1s. If the current position of the animal after the 6s-sound differed from the original position, no power would be energized.

The following experimental parameters of the shuttle box were set, namely, conditional stimulation type: acoustic stimulation, CS duration: 6s, US duration: 6S, unconditional stimulation intensity: 0.5mA, habitation time: 5 min, training interval: 29s, training times: 50 times per 12h.

#### Multi-type shuttle box training

Multi-type shuttle box training refers to change the training box of a range of shapes for respective training based on the Sound-cued active avoidance training. In the present study, three types of training boxes with a range of shapes were employed, namely, a standard square shuttle box, a shuttle box recessed in the middle, as well as a contraction shuttle box on either side (Fig. 1 A). All the shuttle boxes exhibited similar internal surface area and volume except for their shapes.

**Figure 1.**
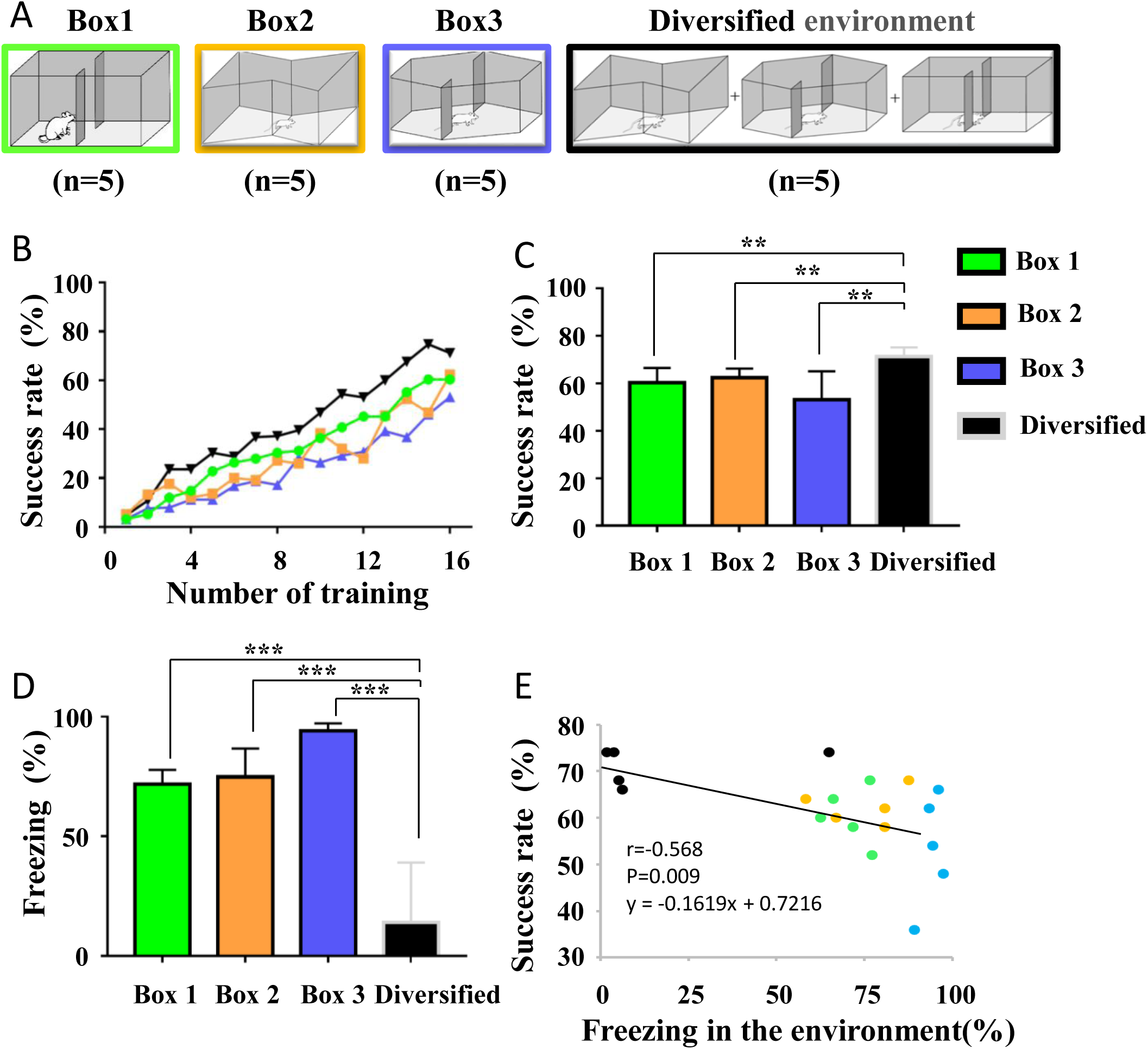
Diversified environment can improve learning in active avoidance. (A) Multi-type shuttle boxes; (B) Success rate of animals trained in four types of environment; (C) The average success rate of last 3 trainings in A; (D) Freezing proportion in four types of environment; (E) Linear regression analysis between success rate and freezing proportion. Green: box 1; orange: box 2; purple: box 3; black: diversified environment. **: p < 0.01; ***: p < 0.001.

#### Determine the extent of the fear of the training context

In the experiment performed in this study, animals trained with the use of various training methods displayed different stress levels of the training environment. A separated behavioral paradigm was adopted to identify the animal’s fear of training environment before or after training.

The animal was placed in the context to be tested. After the animal was acclimated for 1 min, the animal’s behavior was recorded for 5 min, and the ratio of time when the animal was not moving in the context referred to the extent of the fear of training context. The freezing level was scored when no movement (except for respiratory movements) was detected for at least 1s, and the total freezing time during the sound presentation was counted by video analysis.

#### Preconditioning (unconditioned foot-shock)

A paradigm was also set to raise the fear of animals in a context. The animals were placed in the context where raised fear was required. Moreover, the animals were given a continuous electric shock (0.5 mA) for 10 s, which were repeated 10 times at intervals of 50 s.

#### Opening field test (OFT)

OFT was performed as mentioned previously to measure the spontaneous activity of rodents. In brief, the device consisted of a gray square (100 cm to 100 cm to 40 cm). It was split into 25 cm to 25 cm equal squares drawn on the arena board. The test room was dimly lit (the identical light to the rearing environment). A single rat was placed in the center of the floor. Subsequently, after the rat underwent 30 s adaptation process, the number of squares crossed (with the four paws), the number of rears (posture sustained with hind-paws on the floor) were counted manually for 5 min, as well as the time spent on grooming (e.g., washing or mouthing of forelimbs, hind-paws, face, body and genitals). All behavior was recorded with a video camera located 150 cm above the arena as described previously. After each test was performed, the arena was cleaned with 75% alcohol solution.

#### Forced swimming test (FST)

The trained rats were placed in a container filled with water. The container was a transparent cube, 25 cm in length, 25 cm in width, and 35 cm in height. The water depth was 25 cm, and the water temperature was 24 degrees. The animals started to count for 5 min. The whole experiment was performed in the same lighting environment to the rearing. The immovable was defined that the entire body had no other movement except for the tiny movements required to keep the animal’s head above the water surface. The percentage of the animal’s immobility time in the 5 min was counted. After the experiment was performed, the animal’s body was wiped with/by a dry towel, and the animal’s body was dried with hot air. Subsequently, the animal was placed in a feeding cage.

#### Elevated plus-maze test (EPM)

The elevated Plus-maze was made of odorless plastic material with a color of black. It covered two relatively open arms of 50cm by 10cm, as well as two relatively closed arms of 50cm by 10 cm by 40 cm. There was an open area of 10cm by 10cm in the center of the maze. The maze was 100 cm in height. A video monitor was set 150cm above the maze.

The experimental animal was placed in the central area of the maze, with the head facing towards the open arms. It is noteworthy that each experimental animal would be arranged in the identical position thereafter.

In the meantime, the camera monitor was opened to record the time that the experimental animals accessed into the arms with the open and closed arms within 5 min. During the experiment, the experimenter should be 1 m away from the maze. After the recording was finished, the experimental animals were returned to the breeding cage. The maze was also cleaned and wiped with 75% alcohol to avoid being affected by the animal odor on subsequent experimental animals. The proportion of the time which each animal staying in the open arm was calculated after the experiment. The percentage of open arm retention time was expressed as:

Percentage of open arm retention time = open arm retention time / (open arm retention time + closed arm retention time).

#### Light-dark transition test

The experiments were performed in the experiment box applied here following the previous procedures [31]. The device consisted of a box (30 × 30 × 70 cm), and the cage was split into two chambers of equal size by a partition with a door. One chamber was composed of white plastic walls; it was brightly illuminated (390 lux) by lamps above the ceiling. The other chamber showed black plastic walls; it was dark colored (2 lux). Both rooms had grey plastic floors. The mice were placed in a dark room and subsequently moved freely between the two rooms with the door open for 5 min. After testing, the proportion of time spent by mice in the bright room was calculated.

## Results

### Diversified environment can improve learning in active avoidance

The active avoidance paradigm in classic sound cued two-way shuttle boxes was first established in accordance with the existing study[27]. It was showed that adult rats exhibited similar learning capabilities in three boxes (Fig. 1B & C), revealing that the designs of three boxes did not noticeably change the learning performance of animals. In contrast to animals trained in single shuttle box, animals trained in diversified environments displayed a steeper learning curve (Fig. 1B) and had a higher success rate (Fig. 1C). Twelve hours after the last training was conducted, animals tested in single shuttle box showed a high level of freezing behavior (Box 1: 71.67 ± 2.51%; Box 2, 74.83 ± 5.31%; Box 3, 94.07 ± 1.40%) when no sound cue or foot shock was delivered. Accordingly, a huge stress the animal felt in training box was suggested. Meanwhile, animals trained in diversified environments displayed low levels of freezing behavior (14.05 ± 10.21%, n = 5) when left in the most recent training box, which were statistically different from those of the other groups (Fig. 1D). It was therefore revealed that when trained in diversified environments, animals could maintain a low stress level in training box. According to the results of linear regression analysis, the level of animal stress was negatively correlated with behavioral performance (R = 0.568, p = 0.009) (Fig. 1E). As suggested from the mentioned, diversified environments could improve learning ability in active avoidance by lowering animal stress in training boxes.

### Stressful precondition lower performance in active avoidance learning

To delve into the role of environmental stress in active avoidance learning, precondition was employed to manipulate animal stress in a certain training box (Fig. 2A). A one-time unconditioned foot-shock (0.5 mA for 10 sec, 50 sec interval, 10 repetitions) acted as the stressful precondition 12 h before the first training session of active avoidance learning. Two groups of animals were used for the following study. For the group exhibiting high stress level, the animals were trained in the same box as preconditioning. In terms of the group displaying low stress level, animals received the training in a novel box. A significant level of environmental stress was identified when animals were left in preconditioned box in contrast to those moved to a novel training box (high stress: 58.48 ± 8.82%; low stress: 20.00 ± 8.00%) (Fig. 2C). It was therefore demonstrated that preconditioning with foot-shock could successfully elevate animal stress in a certain training box. As expected, better learning performance was found in the low stress group as compared with the high stress group (Fig. 2B). After being trained, animals in both groups exhibited high levels of stress in their training boxes (Fig. 2C), whereas only animals in the low stress group showed significant increase of stress (Fig. 2C). According to these results, the training protocol could elevate animal stress in the training box, whereas the level of stress was already saturated right after preconditioning in high stress group.

**Figure 2.**
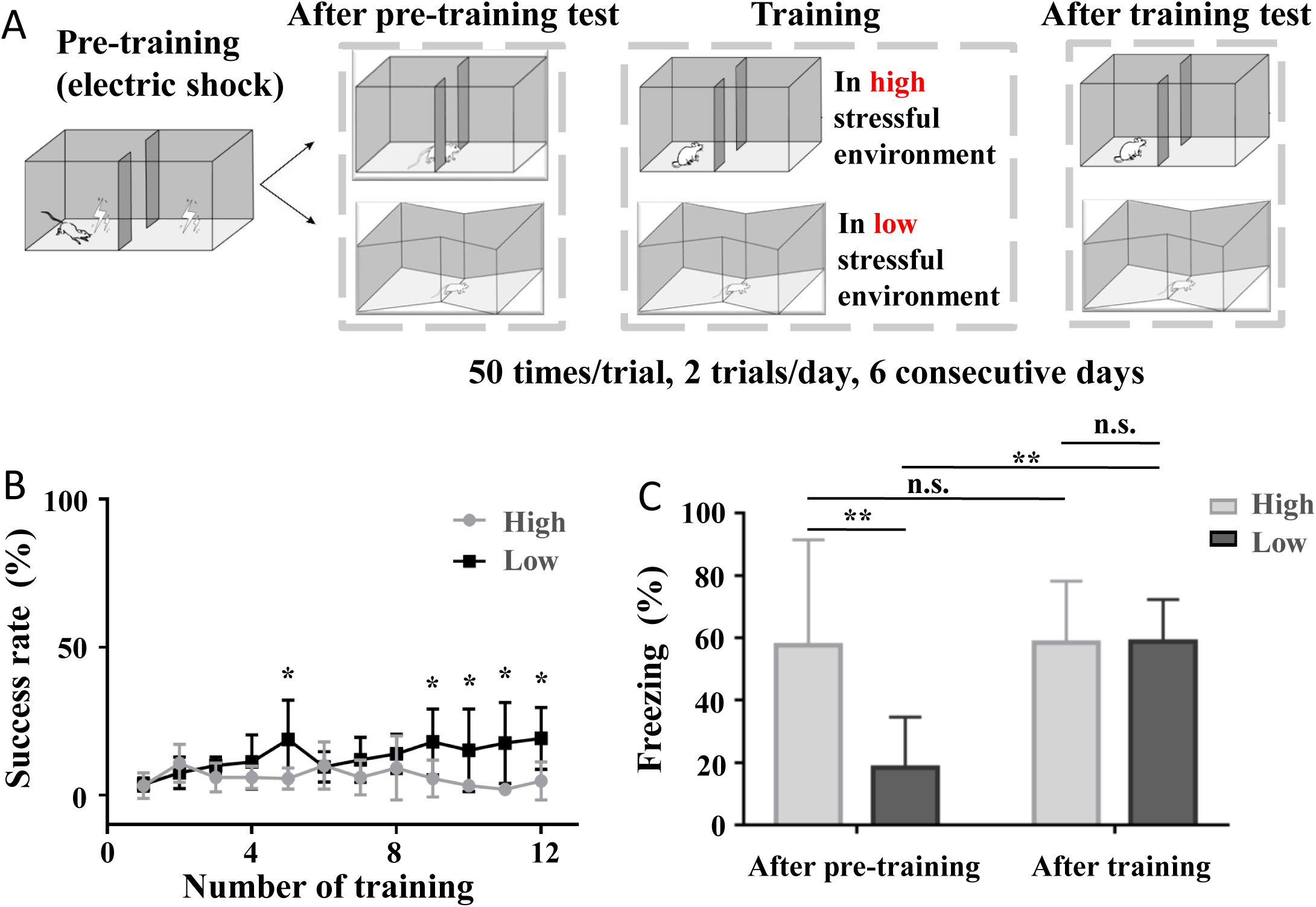
Stressful precondition lower performance in active avoidance learning. (A) Training flow chart; (B) Success rate of animals trained in high stressful environment (in the same box) and low stressful environment (in a novel box); (C) Freezing proportion of pre-training and after training in high stressful environment or low stressful environment. Grey: in high stressful environment; dark: in high stressful environment; *: p < 0.05; **: p < 0.01; n.s.: no significant difference.

To further examine the depression/anxiety level of animals in two groups, elevated plus maze (EPM, Fig. 3A & B), force swimming test (Fig. 3C), light-dark box test (Fig. 3D), and open field test (Fig. 3E-G) were performed in this study. No significant difference was found between high stress group and low stress group after training, demonstrating that distinct environmental stress was the major difference and could be the cause of learning difference between two groups.

**Figure 3.**
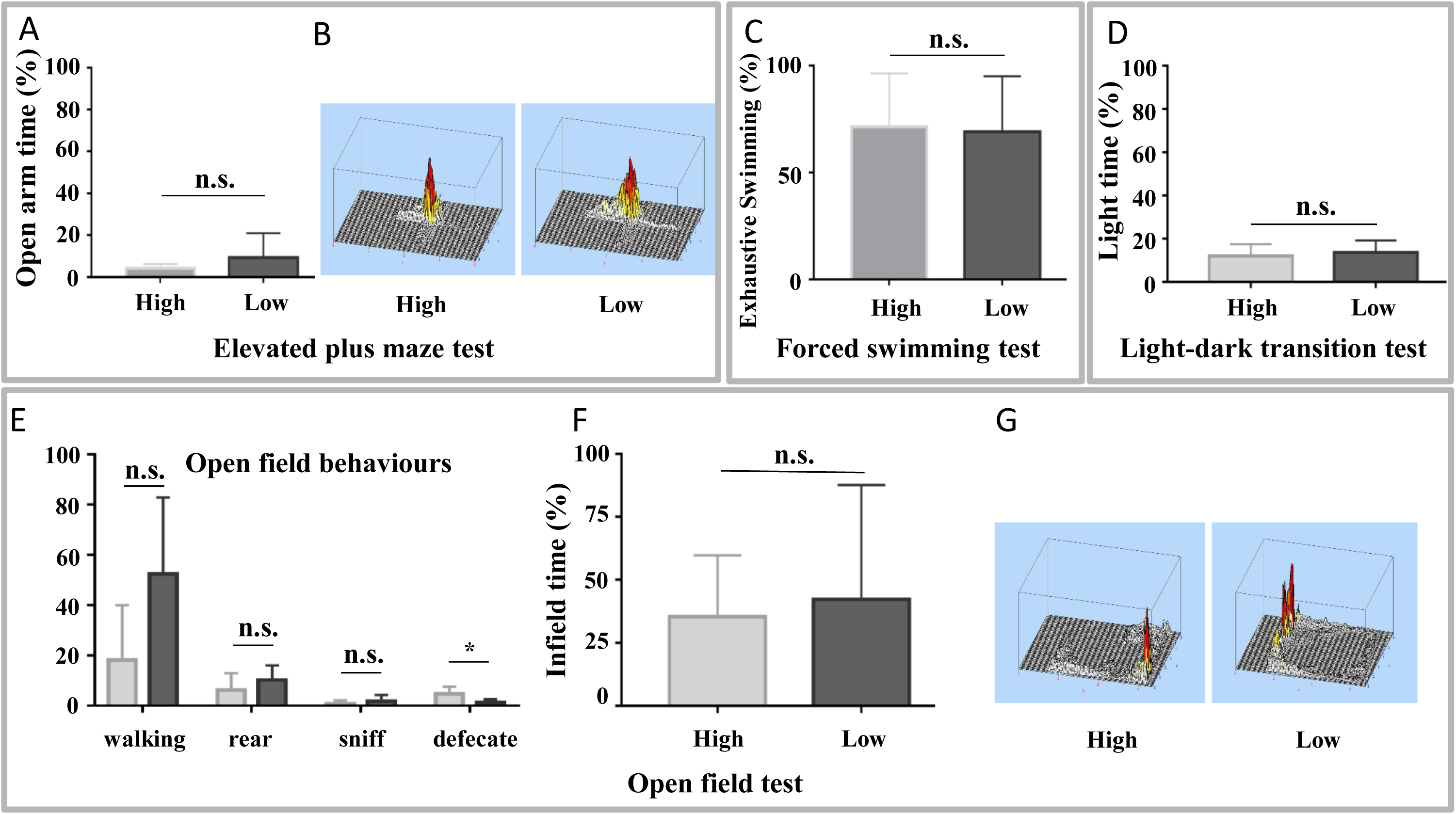
Depression/anxiety level examination of animals trained in high stressful environment and low stressful environment. (A) Proportion of open arm time in elevated plus maze test; (B) 3D activity distribution of elevated plus maze test; (C) Proportion of exhaustive swimming time in forced swimming test; (D) Proportion of the light time in light-dark transition test; (E) Open field behaviours (walking, rear, sniff, defecate) in open field test; (F) Proportion of the infield time in open field test; (G) 3D activity distribution of open field test. Grey: in high stressful environment; dark: in high stressful environment; *: p < 0.05; n.s.: no significant difference.

### Learning impairment caused by stressful context can be alleviated by training in diversified training environment

Because diversified training environments have a positive impact in reducing environment stress (Fig. 1), it is possible that diversified training environments could also improve learning impairment attributed to environmental stress. Preconditioned animals were trained in single shuttle box or diversified environments, respectively (Fig. 4A). Better learning scores (Fig. 4B) and lower stress level (Fig. 4C) were identified in animals trained in multiple shuttle boxes. Note that the stress level did not significantly increase after training (Fig. 4C) when preconditioned animals were trained in diversified environments. Compared with the elevated stress level shown in Fig. 2C, this result suggested that diversified environments can effectively inhibit environmental stress from increasing during training. Meanwhile, similar level of depression/anxiety was found between two groups (Fig. 5A-G). These results are in accordance with previous results and supports our hypothesis that diversified training environments can reduce negative influence attributed to environmental stress.

**Figure 4.**
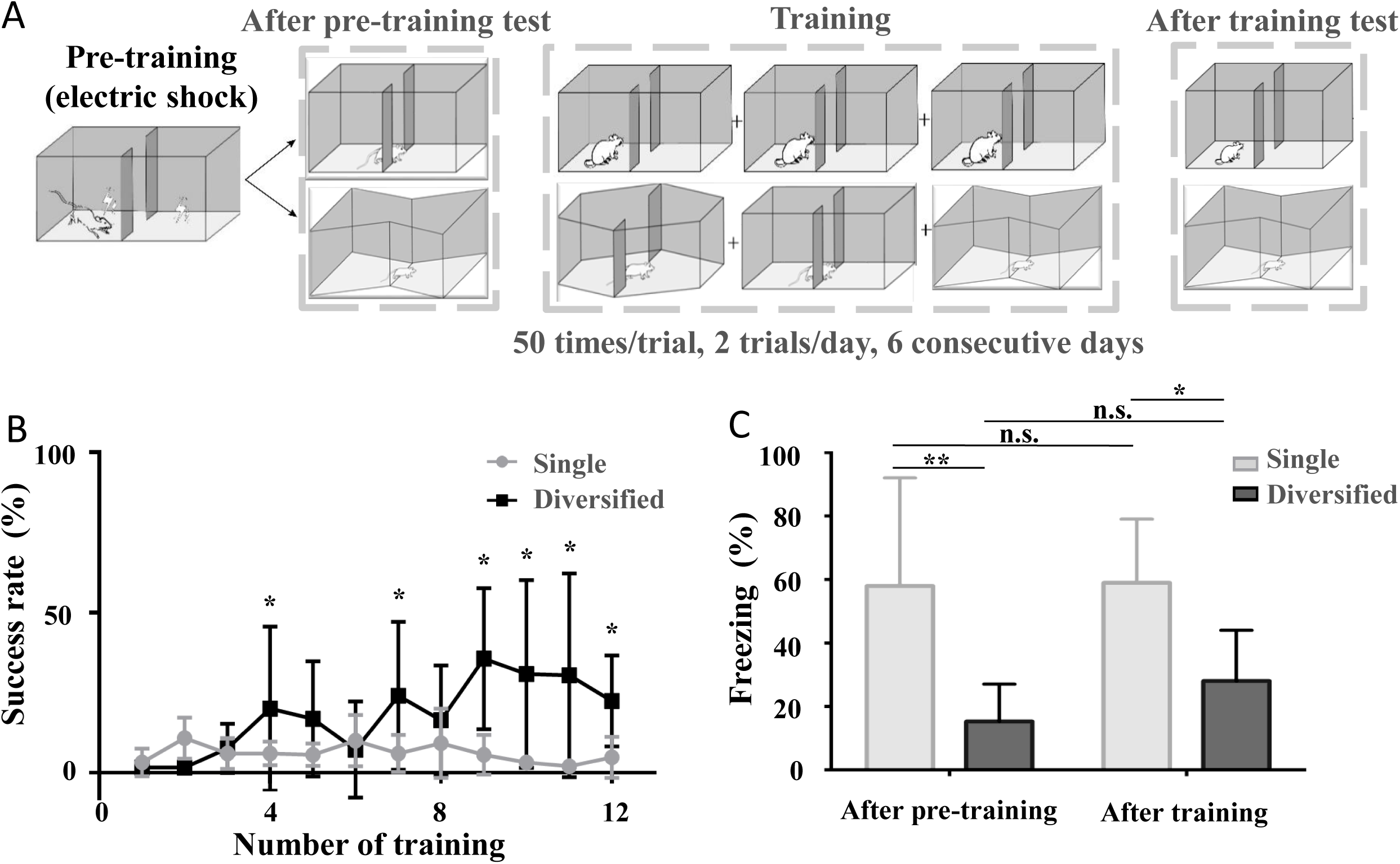
Learning impairment caused by stressful context can be alleviated by training in diversified training environment. (A) Training flow chart; (B) Success rate of animals trained in single environment and diversified environment; (C) Freezing proportion of pre-training and after training in single environment or diversified environment. Grey: in single environment; dark: in diversified environment; *: p < 0.05; **: p < 0.01; n.s.: no significant difference.

**Figure 5.**
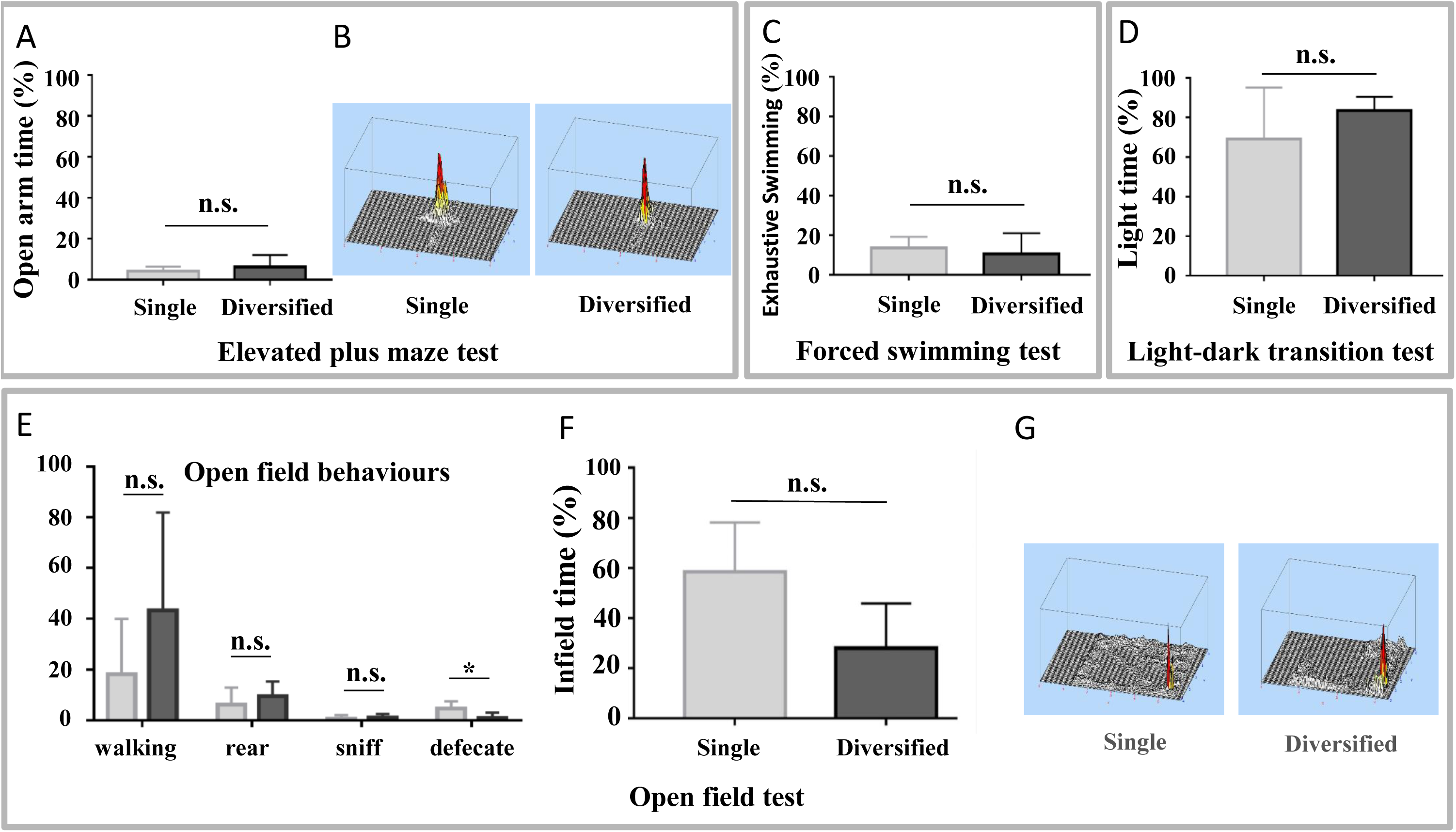
Depression/anxiety level examination of animals trained in single environment and diversified environment. (A) Proportion of open arm time in elevated plus maze test; (B) 3D activity distribution of elevated plus maze test; (C) Proportion of exhaustive swimming time in forced swimming test; (D) Proportion of the light time in light-dark transition test; (E) Open field behaviours (walking, rear, sniff, defecate) in open field test; (F) Proportion of the infield time in open field test; (G) 3D activity distribution of open field test. Grey: in single environment; dark: in diversified environment; *: p < 0.05; n.s.: no significant difference.

## Discussion

From the ecological perspective, an enriched breeding/living environment provides animals with greater environmental tolerance. Numerous studies reported that animals under enriched living environments could exhibit better performance on learning tasks and exhibit higher levels of exploration activity with lower anxiety[12, 14, 20, 32]. A considerable previous research have shown that enhanced environmental complexity could cause behavioral changes related to various variations in the brain[13, 19, 24]. Enriched environments are critical to the nervous system of juvenile and adult animals; they are likely to help reverse cognitive and emotional disorders. For mammals, the environment where they grow up noticeably impacted their negative emotional and neuronal development [33-35]. As revealed from the results here, for adult rats, simply increasing the diversity of training environments could effectively elevate the learning efficiency of animals without the need to enrich breeding environments. Accordingly, a novel strategy to elevate learning efficiency and mitigate environmental stress was also presented.

Previous studies have shown that strengthened synaptic connection was found in hippocampus[35]and cortex[33]of animals housed in enriched environments and could be the neural mechanism responsible for the enhanced learning performance. Moreover, activation of prefrontal cortex in rats could enhance active avoidance performance and reduce freezing behavior to neutral conditioned stimuli by inhibiting neuron activities in amygdala[26]. Though the neural mechanism has not been explored in this study, the cellular plasticity and circuitry modulation identified in existing studies could also contribute to increased learning in diversified environments.

Another noteworthy finding achieved here was about the reduced environmental stress caused by diversified environment. The amygdala has been known as the center of fear-related memories [22]. In human studies, increased activity was found in the amygdala of patients with depression [36, 37]. Further studies have reported that depression could be relieved by suppressing the amygdala[30, 38]. In the meantime, researchers also suggested that living in a stressful urban environment could also enhance amygdala activity[1, 6, 14]. As revealed from the results here, by diversifying the environment, the stress generated during the training process could be lowered. The results here indicated a possibility that diversified learning environment may inhibit the activity level of the amygdala and can facilitate the treatment of mental disorders attributed to negative emotions. By diversifying the environment, the environment stress could be reduced during the learning process, so the subject could focus more on the learning task itself. The results here might also explain the reason why exchange students and field training of professional teams can enhance learning efficiency.

## Acknowledgments

National Natural Science Foundation of China (No. 31600848), Chongqing University of Science and Technology Research Foundation (No. CK2016Z02).

